# Graphsite: Ligand-binding site classification using Deep Graph Neural Network

**DOI:** 10.1101/2021.12.06.471420

**Authors:** Wentao Shi, Manali Singha, Limeng Pu, J. Ramanujam, Michal Brylinski

## Abstract

Binding sites are concave surfaces on proteins that bind to small molecules called ligands. Types of molecules that bind to the protein determine its biological function. Meanwhile, the binding process between small molecules and the protein is also crucial to various biological functionalities. Therefore, identifying and classifying such binding sites would enormously contribute to biomedical applications such as drug repurposing. Deep learning is a modern artificial intelligence technology. It utilizes deep neural networks to handle complex tasks such as image classification and language translation. Previous work has proven the capability of deep learning models handle binding sites wherein the binding sites are represented as pixels or voxels. Graph neural networks (GNNs) are deep learning models that operate on graphs. GNNs are promising for handling binding sites related tasks - provided there is an adequate graph representation to model the binding sties. In this communication, we describe a GNN-based computational method, GraphSite, that utilizes a novel graph representation of ligand-binding sites. A state-of-the-art GNN model is trained to capture the intrinsic characteristics of these binding sites and classify them. Our model generalizes well to unseen data and achieves test accuracy of 81.28% on classifying 14 binding site classes.

## 1 Introduction

Interactions of proteins with other molecules like peptides, neurotransmitters, nucleic acids, hormones, metabolites, and other proteins have vital part in understanding the biological functions. Proteins are basic building blocks and responsible for carrying out all biological functions in cellular environment. So, identification of interaction between proteins and small molecules is crucial to understand how proteins regulate different functions in a living cell [2]. The ligand binding site (also referred to as pocket) is a groove or cavity in a protein where the small molecules or ligands can bind through interactions with amino acids at that site [27]. Identification of off-targets binding can help scientists to repurpose the existing drugs to cure some rare orphan diseases for which we do not have functional drugs available. So, binding site prediction approaches can be used to find cures for rare diseases [10]. Therefore, binding site prediction in structural biology is vitally important in the field of drug discovery and it can help predict the novel drug targets. There are several available algorithms which can identify the ligand binding sites on target protein structures such as eFindSite [7], Fpocket [20], and FTSite [25] etc. Besides that, the ligand binding on protein depends on numerous factors of binding site. So, there are various methods which account these factors such as conformational dynamics [1], druggability [16] and amino acid compositions [31] of binding sites on target proteins. However, all these methods do not account for the classification of binding sites depending on types of ligands.

Deep learning is an emerging machine learning technique. Deep learning-based models employ various styles of multi-layer artificial neural networks to learn from data and make predictions. Deep learning has achieved significant progress in computer vision applications such as object detection [13], face recognition [28], and human pose estimation [35]. One of the keys to the success of those applications is the convolutional neural network (CNN), which can learn hierarchical latent features from Euclidean data (2D- and 3D images) by utilizing local trainable filters [3]. Such methodologies in computer vision have inspired new works in structural biology in recent years. DeepDrug3D [26] achieves state-of-the-art binding site classification performance by representing the binding sites as 3D images and deploying a 3D-CNN. DeeplyTough [30], which uses similar pocket representation as DeepDrug3D, implements pocket-matching. DeepSite [15] is a binding site predictor that also forms similar 3D representations of pockets by computing atomic-based pharmacophoric properties for each voxel. Other than 3D representations, BionoiNet [29] projects pockets to 2D images that encode chemical properties, and a 2D-CNN is trained to perform classification.

Graph neural network (GNN) is another category of deep learning model that operates on graphs which are non-Euclidean data. Over the recent years, GNNs have demonstrated encouraging performance on applications such as text classification [12, 18] and traffic prediction [22]. As for the field of chemistry and biochemistry, GNNs are proven to be promising for a variety of applications including predicting quantum property of an organic molecule [9], generating molecular fingerprints [5], predicting protein interface [8], and predicting drug-target interaction [23]. These works are based on the idea that molecular structures can be naturally interpreted as graphs. A typical example is the Lewis structure of molecules where the atoms are treated as nodes and the chemical bonds are the undirected edges that connect nodes.

In this communication, we describe a framework based on GNN to classify ligand-binding sites. A novel graph representation of binding sites is developed and a GNN classifier is then trained to classify a pocket dataset of 14 classes. Comparing with the methods that convert pockets to Euclidean data, the process of converting to graphs is fast and lossless. So, the graphs can be generated on-the-fly and the users only need to provide standard text files as input. Our implementation achieves state-of-the-art performance and the followed case studies show that our model learns the underlying pattern of different kinds of binding pockets.

## 2 Materials and Methods

### 2.1 Graph representation of biding sites

The pockets are transformed to graphs as the input of the classifier. The nodes of the graph are the atoms, and an undirected edge is formed between two atoms if the distance between them is less than or equal to 4.5 Å. We crafted 11 node features, 7 of them are spatial features, and the other 4 features are chemical features. The spatial features are used to define the shape of the binding pocket, which are the Cartesian coordinates (x, y, z) of the atoms, the spherical coordinates (r, theta, gamma) of the atoms, and the solvent accessible surface area (SASA). We adopt the chemical features described in Bionoi [6], which are charge, hydrophobicity, binding probability and sequence entropy. Fig 1 illustrates part of the graph representation of a binding pocket. As can be seen in Fig 1B, each atom is connected to all the neighboring atoms withing the radius of 4.5 Å. To distinguish the chemical bond-edges from the others, we set the number of chemical bonds as the edge attribute. The edges with no chemical bonds have 0 as their attributes and the edges on aromatic rings have 1.5 as their attributes.

**Figure 1.**
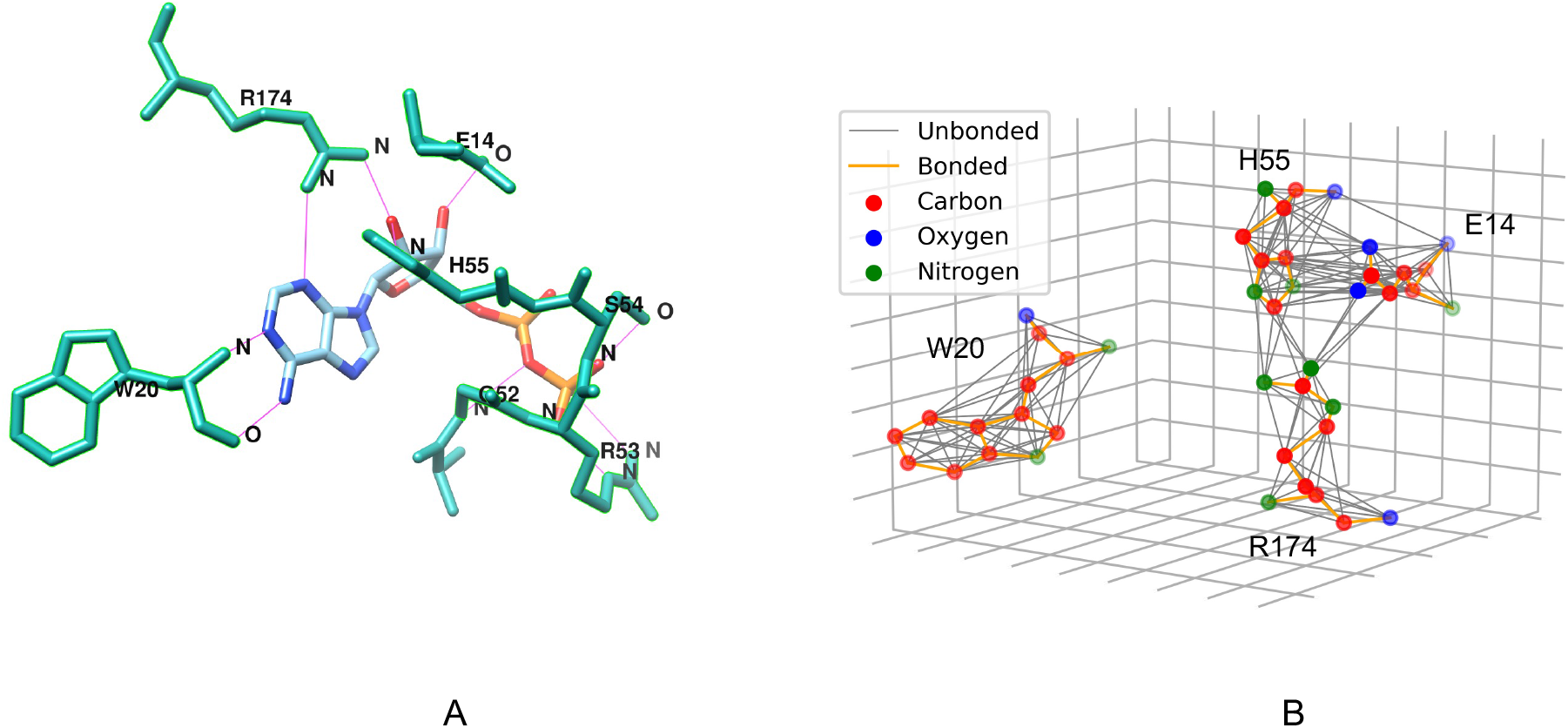
Molecular structure and graph representation of a binding site. (A) The residues that interact with the ligand of pocket 5×06F00. (B) The graph representation of 4 residues in A. Any atom pair that has distance less than or equal to 4.5 Å is connected

### 2.2 Graph neural network

With the graph representation of the binding pockets, the binding site classification problem essentially becomes a graph classification problem. A general graph classification framework that uses GNN can be divided into three stages: message passing, graph readout, and classification. In addition to these three stages, our model utilizes jumping knowledge connections [37] to let the model select information for each node from different layers. Fig 2 illustrates the overall architecture of the GraphSite classifier. As can be seen in Fig 2, the main body of the classifier is an embedding network which contains the message passing layers, the jumping knowledge connections, and a global pooling layer which performs the graph readout. The node features of input graph are updated by the message passing layers. The outputs of all layers are processed by a max pooling layer that performs a feature-wise max pooling; the intuition behind this it to let the model to learn the proper number of layers for each individual node; this technology is known as the jumping knowledge [37]. The max pooling layer is followed by a global pooling layer, which reduces the dimension of node feature from *n × d* to d where *n* is the number of nodes and *d* is dimension of the node feature. The output of the global pooling layer is a fixed-size vector, and it is followed by fully connected layers to generate the final classification results.

**Figure 2.**
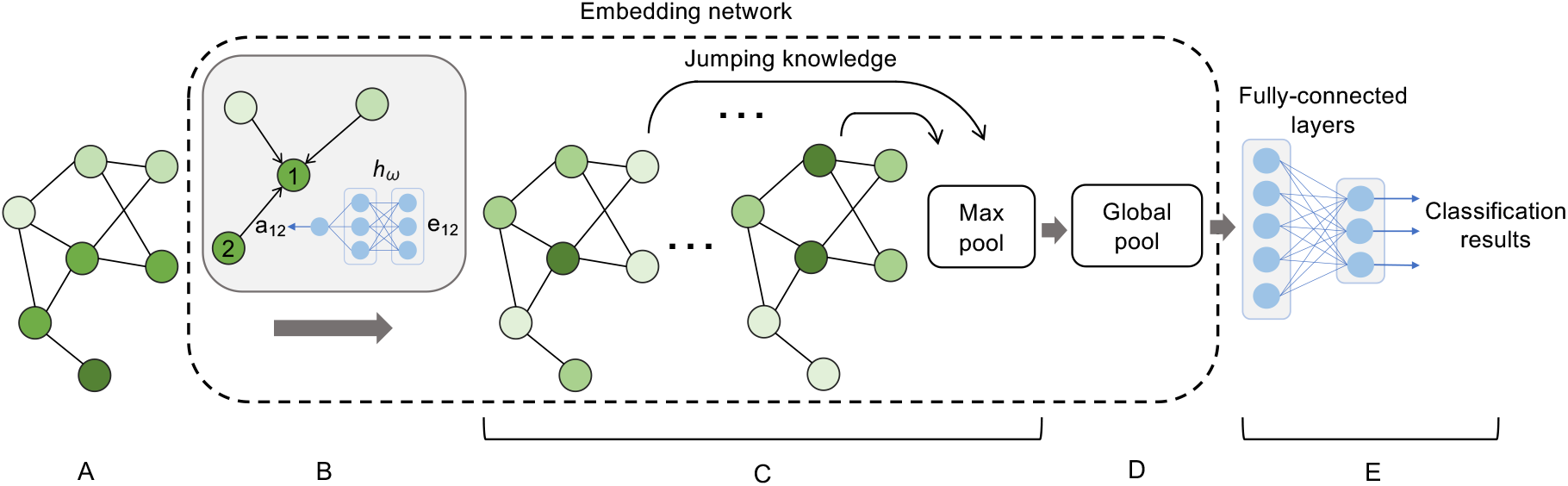
The architecture of GraphSite classifier. (A) The input graph of a binding site. (B) A neural network that computes the weight of message using the edge attribute as input. (C) The message passing layers with Jumping Knowledge connections. (D) The global pooling layer which is the Set2Set model. (E) The fully connected layers that generate classification results.

#### 2.2.1 Message passing

The message passing layers of GNNs update the node features by propagating information along edges. From the perspective of each node, the information of its neighborhood is aggregated, and the updated node features can reveal informative local patterns. As described in [41], most of the message passing layers fall into the general form of neighborhood aggregation:

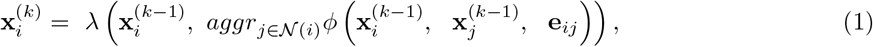

where *ϕ* is a differentiable function that generates the message, aggr is a permutation-invariant function (such as sum or max) that aggregates the messages, and *λ* is a differentiable function such as a multi-layer perceptron (MLP). 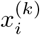 is the output node feature of node *i* of layer 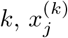 represents its neighbor nodes, and *e*_*ij*_ is the edge attribute. To exploit both node features and edge feature of the binding site graph, we develop a message passing layer which also falls into the general form described by Equation 1, which is called neural weighted message (NWM):

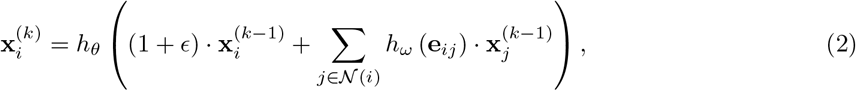

where *h*_*ω*_ is an MLP that takes the edge attribute as input and outputs a scaler as the weight of the message, which is simply node feature *j*; *ϵ* is learnable scalar; *h*_*θ*_ is another MLP that updates the aggregated information. Note that the edge attributes are not updated during training, and they are the same for all the layers. Fig 2B demonstrates an example of NWM: *h*_*ω*_ takes the edge attribute **e**_12_ as input, generating **a**_12_ as the weight of message propagating from node 2 to node 1.

The NWM message passing rule can be regarded as an extension of the graph isomorphism network (GIN) [36]. GIN is an expressive message passing model that is as powerful as the Weisfeiler-Lehman test in distinguishing graph structures; we replace its sum aggregator with sum of weighted messages where the weights are generated by a neural network *h*_*ω*_. From another perspective, the NWM layer belongs to the Message Passing Neural Network (MPNN) family [9]. The gated graph neural network (GG-NN) is an MPNN family member and its message is formed by 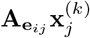, where 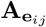 is a square transformation matrix generated by an MLP which takes the edge attribute **e**_*ij*_ as input; if we put a restriction on the matrix 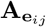, such that it is a diagonal matrix and all elements on the diagonal are equal, the GG-NN module becomes NWM. In fact, the neural message of GG-NN was one of our first design choices. In our experiments, we found that regularizing GG-NN to NWM could help mitigate overfitting and NWM is more computationally efficient. Therefore, we take NMM as our final design choice.

Finally, inspired by the idea that multiple aggregators can improve the expressiveness of GNNs [4], we extend a single-channel NWM layer described by Equation 2 to a multi-channel NWM layer by concatenating the outputs of multiple aggregators:

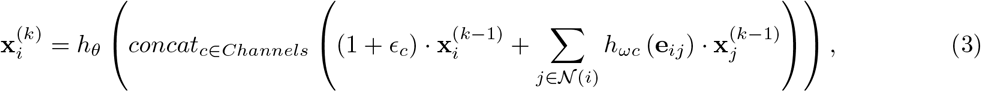

where each pair of *ϵ*_*c*_ and *h*_*ωc*_ represents an aggregator learned as channel c, and C denotes the set of channels. The aggregated node features are concatenated in their last dimension such that the concatenated node features have the shape of *n* by *d ×* |*C*| where *d* is the dimension of node feature. Accordingly, the update neural network *h*_*θ*_ now also acts as a reduction function that reduces the size of node feature from *d ×* |*C*| to *d*. Intuitively, the concatenation of multiple aggregators is analogous to using multiple filters in CNN: each aggregator corresponds to a filter, and the concatenated output corresponds to the output feature maps in a convolution layer in CNN.

#### 2.2.2 Graph readout

The graph readout function reduces the size of graph to one node. This function should regard the features of the nodes as a set, because there is no order among the nodes. i.e., the graph readout function should be permutation invariant. The Set2Set [34] model is a global pooling function to perform graph readout. Set2Set can generate fixed-sized embeddings for sets with various sizes, and it bears the property of permutation invariance. It computes the global representation of the set by leveraging the attention mechanism. Basically, a Long short-term memory (LSTM) [14] neural network recurrently updates a global hidden state of the input set; during the recurrent process, the global hidden state is used to compute the attention associated with each element in the set, and these attentions are in turn used to update the global hidden state. After several such steps, the global graph representation is formed by concatenating the global hidden state generated by the LSTM and the weighted sum of the elements in the set.

#### 2.2.3 Loss function

Instead of the cross-entropy loss, the focal loss [24] is used instead. As will be described in later section, the dataset has imbalanced classes. Some classes such as ATP have much more data points than others. Therefore, most of the data in a mini batch will come from these major classes and the cross-entropy loss will be dominated by them. To mitigate this problem, the focal loss adds a damping factor (1−*p*_*t*_)^*γ*^ to the cross-entropy loss:

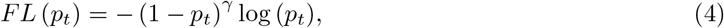

where *p*_*t*_ is the predicted probability generated by the Softmax, and *γ* ≥ 0 is a tunable hyper-parameter. With this damping factor, the dominating confident predictions with high probabilities will be suppressed and the predictions with low probabilities will have higher weights. As a result, the dominated minority classes with low probabilities will have higher weights, and the problem of imbalanced classes is improved.

### 2.3 Dataset

The dataset is generated by clustering the pockets according to their Tanimoto coefficients of the ligands, because similar ligands bind to similar pockets. Note that identical pockets are removed from the dataset. During our experiments, we found that some of the pocket clusters generated by this algorithm are highly similar. We manually identified the type of ligands that bind to each class and found that due to the large Tanimoto distance threshold in clustering, pockets from the same family are divided into different clusters. For example, as illustrated in Fig 4, cluster 0 and cluster 9 are ATP-like pockets, and cluster 3 and cluster 8 are both glucopyranose-related pockets. As a result, 30 largest clusters are selected, and they are merged into 14 classes. The labels of the 14 classes are shown in Table 1.

**Table 1.** The 14 labels of binding sites in the dataset.

| Class | Label |
| --- | --- |
| 0 | ATP |
| 1 | Heme |
| 2 | Carbohydrate |
| 3 | Benzene ring |
| 4 | Chlorophyll |
| 5 | Lipid |
| 6 | Essential amino/citric/tartaric acid |
| 7 | S-adenosyl-L-homocysteine |
| 8 | CoenzymeA |
| 9 | Pyridoxal phosphate |
| 10 | Benzoic acid |
| 11 | Flavin mononucleotide |
| 12 | Morpholine ring |
| 13 | Phosphate |

## 3 Results and discussions

In this section, we first discuss the classification performance of GraphSite classifier along with the baseline methods; then some interesting cases from the misclassified binding pockets are selected as case studies. Finally, we test our model on unseen data which are uploaded to PDB after the curation of our dataset.

### 3.1 Classification performance

Two GNN-based methods are evaluated: GraphSite and GIN. GIN uses a sum aggregator, so the edge attributes are ignored. The purpose of having GIN as a baseline is to demonstrate the improvement of NWM which utilizes edge attributes. The configurations of GraphSite and GIN are identical except the architecture of GNN layers. Both models are trained with the Adam [17] optimizer for 200 epochs and identical learning rate schedulers are used to half the learning rate at plateau. 25 experiments are conducted for each model. In each experiment, each class is randomly divided into a training set and a testing set with different random seeds. After training, the medium accuracies among the 25 experiments on test set are used to evaluate the classification performance. In addition, docking and pocket matching are also tested on the same classification task. We select SMINA [19], which is based on Auto-dock Vina [32] as the docking tool. As for pocket matching, G-LoSA [21] is selected. Since there is no training required for docking and pocket matching, the accuracies over the entire dataset are reported. For docking, a label ligand is chosen manually for each class. For each prediction, the docking score of the pocket is evaluated against all 14 label ligands, and the predicted class is the ligand with best docking score. Pocket matching is conducted in a similar way: a label pocket is chosen for each class, and the predicted class is the label pocket that has best matching score with the pocket to predict. Table 2 shows the classification performance. As shown in Table 2, GraphSite achieves the best overall classification accuracy of 81.28%, along with a weighted F1-score of 81.66%. The accuracy is of GIN is 75.09%, and its weighted F1-score is 74.35%. The accuracy gain of 6.59% comes from replacing the GNN layers of GIN into multi-channel NWM layers. On the other hand, docking and pocket matching are not working. The reasons can be multifold. First, using one fixed ligand/pocket for each class will decrease the classification performance because they are not necessarily the “golden answer” for each particular pocket. Second, the amount of computation required for docking and pocket matching makes it impractical to run these algorithms exhaustively to maximize the classification accuracy.

**Table 2.** Classification performance.

| Model | Accuracy | Weighted precision | Weighted recall | Weighted F1-score |
| --- | --- | --- | --- | --- |
| Graphsite classifier | 81.28% | 82.33% | 81.28% | 81.66% |
| GIN classifier | 75.09% | 74.26% | 75.09% | 74.35% |
| SMINA | 16.71% | 43.45% | 16.71% | 16.10% |
| G-LoSA | 14.76% | 34.41% | 14.76% | 15.89% |

Fig 3 illustrates the confusion matrix on the test set of our model. As can be seen in Fig 3, each number on the diagonal is a recall of a class; most of the classes are classified well except class 12 and class 13. Class 12 contains morpholine rings, 17% of morpholine rings are misclassified as ATP, and 21% of morpholine rings are misclassified as carbohydrates. Class 13 contains phosphate pockets, 26% of which are misclassified as essential amino acids. The first reason for this is that the support of these two classes in the dataset is low: only 1.77% binding pockets are morpholine rings and only 1.61% binding pockets are phosphate. During training, more gradients will be generated for the majority classes and the model will learn more from the majority classes; applying the Focal Loss only mitigate this problem but cannot fix it completely. The second reason is that, in some of the cases, the binding moiety of the ligand is similar to other types of ligands. For example, the binding moiety of some morpholine rings are highly similar to ATP and carbohydrates. Therefore, the model is in fact making correct predictions about the binding pockets for these cases.

**Figure 3.**
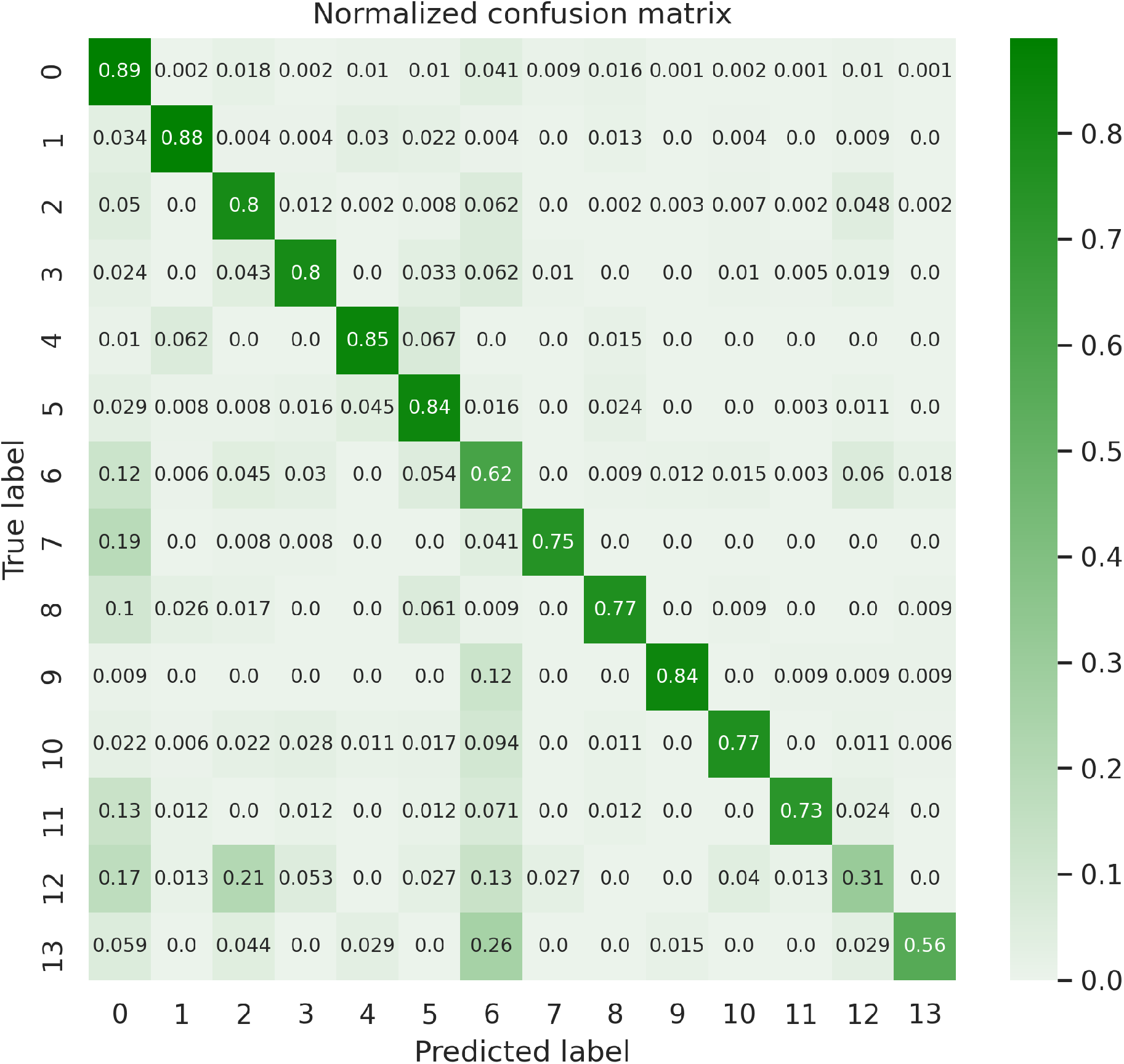
Confusion matrix of the classification result of GraphSite on the test set

## 4 Future work

The performance of Graphsite classifier indictates that the features of ligand-binding pockets are extracted effectively from their graph representations. So, it is possible to extend the settings in this project into other deep learning applications, such as metric learning and generative modeling. In the next chapter, we describe a generative model based on Graphsite for drug discovery. Here, we explore a metric learning model with a Siamese architecture [11] based on Graphsite. After training, the Siamese network can generate embeddings of binding pockets for visualization and other machine learning applications. As can be seen in Fig 4, the embedding network described previously takes a pair of graphs as input and generate two graph embeddings; these embeddings are input of the contrastive loss [11]:

**Figure 4.**
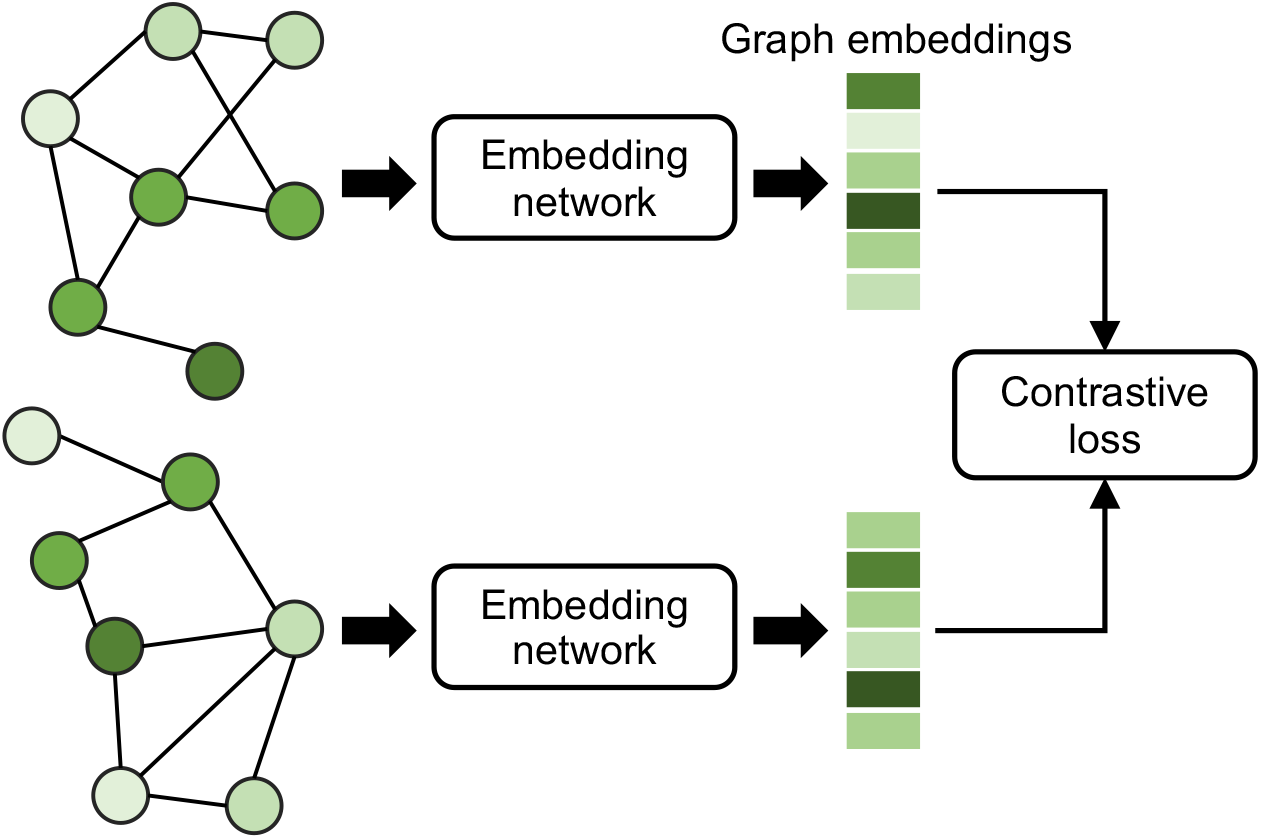
The Siamese-GraphSite architecture. This architecture takes a pair a of graph data as input and it is optimized according to the contrastive loss such that graphs come from the same class are close to each other and graphs from different classes are pushed away from each other.

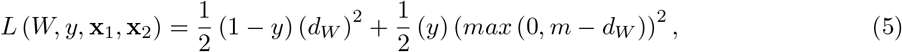

where y is the label of a graph pair that 0 means a similar pair and 1 means a dissimilar pair; **x**_1_ and **x**_2_ are the input graph pair, *W* parameterizes the embedding network, *d*_*W*_ is the Euclidean distance between the graph embeddings, and *m* > 0 is a margin such that a pair contributes to the loss only if their distance is within this margin. Intuitively, the contrastive loss is trying to train a model such that the embeddings from the same class are close to each other in the Euclidean space, and far away from each other if they belong to different classes. Since the model is optimized to manipulate the embeddings in the Euclidean space, the embeddings are ideal for distance-based applications such as t-SNE [33] visualization and k-nearest neighbors. Figure 5 shows the t-SNE visulization of 8 classes from the dataset. As can be seen, similar pockets are clustered together, and dissimilar pockets are separated away from each other, which indicates that the graph Siamese model has learned effective embeddings for the binding pockets. However, as the number of classes increases, the performance of the model decreases significantly in our experiment. We list improving the performance of this metric learning model as one of the future works of Graphsite.

**Figure 5.**
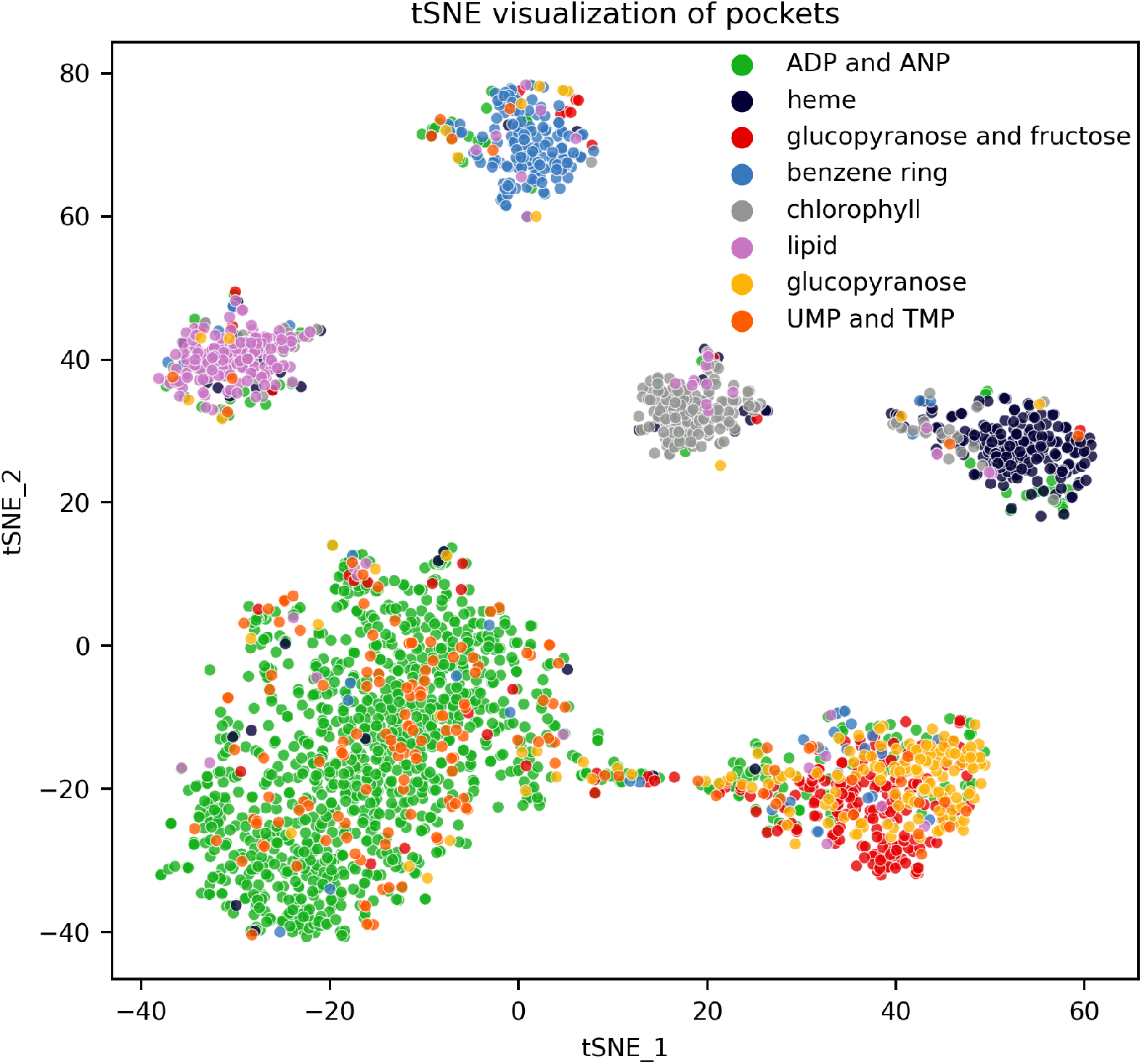
t-SNE visualization of embeddings of selected clusters generated by the Siamese-GraphSite model.

## 5 Conclusion

In this communication, we describe GraphSite, a method to classify ligand-binding sites by modeling ligand-binding sites as graphs and utilizing a GNN as the classifier. The trained model is able to capture informative features of binding pockets, yielding state-of-the-art classification performance. The case studies show that GraphSite successfully classified the binding sites independently of their ligands. Our model is able to make meaningful prediction despite the noise in the dataset caused by the discrepancy between the ligand and its binding moiety. There are several potential ways to improve or extend GraphSite. First, compiling larger datasets with more classes will help training a more power model. Second, exploring more meaningful node features of the binding site graph may also improve the classification performance. Third, GraphSite can be extended to other deep learning-based applications that involve binding sites. For example, it is possible to train a graph autoencoder to generate latent embeddings of binding sites. Another potential application is to build a model to predict drug-target interactions where the GNN layers of GraphSite can be used as the feature extractor of binding sites.

## 6 Supporting Information

- Graphsite is open-sourced and available at https://github.com/shiwentao00/Graphsite.
- The classifier implementation is open-sourced and available at https://github.com/shiwentao00/Graphsite-classifier.

## References

1. M. Araki, H. Iwata, B. Ma, A. Fujita, K. Terayama, Y. Sagae, F. Ono, K. Tsuda, N. Kamiya, and Y. Okuno. Improving the accuracy of protein-ligand binding mode prediction using a molecular dynamics-based pocket generation approach. Journal of computational chemistry, 39(32):2679–2689, 2018.

2. T. L. Blundell. Structure-based drug design. Nature, 384(6604 Suppl):23–26, 1996.

3. M. M. Bronstein, J. Bruna, Y. LeCun, A. Szlam, and P. Vandergheynst. Geometric deep learning: going beyond euclidean data. IEEE Signal Processing Magazine, 34(4):18–42, 2017.

4. G. Corso, L. Cavalleri, D. Beaini, P. Lió, and P. Veličković. Principal neighbourhood aggregation for graph nets. arXiv preprint 2004.05718, 2020.

5. D. Duvenaud, D. Maclaurin, J. Aguilera-Iparraguirre, R. Gómez-Bombarelli, T. Hirzel, A. Aspuru-Guzik, and R. P. Adams. Convolutional networks on graphs for learning molecular fingerprints. arXiv preprint 1509.09292, 2015.

6. J. Feinstein, W. Shi, J. Ramanujam, and M. Brylinski. Bionoi: A voronoi diagram-based representation of ligand-binding sites in proteins for machine learning applications. In Protein-Ligand Interactions and Drug Design, pages 299–312.Springer, 2021.

7. W. P. Feinstein and M. Brylinski. efindsite: Enhanced fingerprint-based virtual screening against predicted ligand binding sites in protein models. Molecular informatics, 33(2):135–150, 2014.

8. A. M. Fout. Protein interface prediction using graph convolutional networks. PhD thesis, Colorado State University, 2017.

9. J. Gilmer, S. S. Schoenholz, P. F. Riley, O. Vinyals, and G. E. Dahl. Neural message passing for quantum chemistry. In International conference on machine learning, pages 1263–1272. PMLR, 2017.

10. R. G. Govindaraj, M. Naderi, M. Singha, J. Lemoine, and M. Brylinski. Large-scale computational drug repositioning to find treatments for rare diseases. NPJ systems biology and applications, 4(1):1–10, 2018.

11. R. Hadsell, S. Chopra, and Y. LeCun. Dimensionality reduction by learning an invariant mapping. In 2006 IEEE Computer Society Conference on Computer Vision and Pattern Recognition (CVPR’06), volume 2, pages 1735–1742. IEEE, 2006.

12. W. L. Hamilton, R. Ying, and J. Leskovec. Inductive representation learning on large graphs. In Proceedings of the 31st International Conference on Neural Information Processing Systems, pages 1025–1035, 2017.

13. K. He, G. Gkioxari, P. Dollár, and R. Girshick. Mask r-cnn. In Proceedings of the IEEE international conference on computer vision, pages 2961–2969, 2017.

14. S. Hochreiter and J. Schmidhuber. Long short-term memory. Neural computation, 9(8):1735–1780, 1997.

15. J. Jiménez, S. Doerr, G. Martínez-Rosell, A. S. Rose, and G. De Fabritiis. Deepsite: protein-binding site predictor using 3d-convolutional neural networks. Bioinformatics, 33(19):3036–3042, 2017.

16. O. Kana and M. Brylinski. Elucidating the druggability of the human proteome with e findsite. Journal of computer-aided molecular design, 33(5):509–519, 2019.

17. D. P. Kingma and J. Ba. Adam: A method for stochastic optimization. arXiv preprint 1412.6980, 2014.

18. T. N. Kipf and M. Welling. Semi-supervised classification with graph convolutional networks. arXiv preprint 1609.02907, 2016.

19. D. R. Koes, M. P. Baumgartner, and C. J. Camacho. Lessons learned in empirical scoring with smina from the csar 2011 benchmarking exercise. Journal of chemical information and modeling, 53(8):1893–1904, 2013.

20. V. Le Guilloux, P. Schmidtke, and P. Tuffery. Fpocket: an open source platform for ligand pocket detection. BMC bioinformatics, 10(1):1–11, 2009.

21. H. S. Lee and W. Im. G-losa: An efficient computational tool for local structure-centric biological studies and drug design. Protein Science, 25(4):865–876, 2016.

22. Y. Li, R. Yu, C. Shahabi, and Y. Liu. Diffusion convolutional recurrent neural network: Data-driven traffic forecasting. arXiv preprint 1707.01926, 2017.

23. J. Lim, S. Ryu, K. Park, Y. J. Choe, J. Ham, and W. Y. Kim. Predicting drug–target interaction using a novel graph neural network with 3d structure-embedded graph representation. Journal of chemical information and modeling, 59(9):3981–3988, 2019.

24. T.-Y. Lin, P. Goyal, R. Girshick, K. He, and P. Dollár. Focal loss for dense object detection. In Proceedings of the IEEE international conference on computer vision, pages 2980–2988, 2017.

25. C.-H. Ngan, D. R. Hall, B. Zerbe, L. E. Grove, D. Kozakov, and S. Vajda. Ftsite: high accuracy detection of ligand binding sites on unbound protein structures. Bioinformatics, 28(2):286–287, 2012.

26. L. Pu, R. G. Govindaraj, J. M. Lemoine, H.-C. Wu, and M. Brylinski. Deepdrug3d: Classification of ligand-binding pockets in proteins with a convolutional neural network. PLoS computational biology, 15(2):e1006718, 2019.

27. D. B. Roche, D. A. Brackenridge, and L. J. McGuffin. Proteins and their interacting partners: An introduction to protein–ligand binding site prediction methods. International journal of molecular sciences, 16(12):29829–29842, 2015.

28. F. Schroff, D. Kalenichenko, and J. Philbin. Facenet: A unified embedding for face recognition and clustering. In Proceedings of the IEEE conference on computer vision and pattern recognition, pages 815–823, 2015.

29. W. Shi, J. M. Lemoine, A.-E.-M. A. Shawky, M. Singha, L. Pu, S. Yang, J. Ramanujam, and M. Brylinski. Bionoinet: ligand-binding site classification with off-the-shelf deep neural network. Bioinformatics, 36(10):3077–3083, 2020.

30. M. Simonovsky and J. Meyers. Deeplytough: learning structural comparison of protein binding sites. Journal of chemical information and modeling, 60(4):2356–2366, 2020.

31. J. Skolnick, M. Gao, A. Roy, B. Srinivasan, and H. Zhou. Implications of the small number of distinct ligand binding pockets in proteins for drug discovery, evolution and biochemical function. Bioorganic & medicinal chemistry letters, 25(6):1163–1170, 2015.

32. O. Trott and A. J. Olson. Autodock vina: improving the speed and accuracy of docking with a new scoring function, efficient optimization, and multithreading. Journal of computational chemistry, 31(2):455–461, 2010.

33. L. Van der Maaten and G. Hinton. Visualizing data using t-sne. Journal of machine learning research, 9(11), 2008.

34. O. Vinyals, S. Bengio, and M. Kudlur. Order matters: Sequence to sequence for sets. arXiv preprint 1511.06391, 2015.

35. S.-E. Wei, V. Ramakrishna, T. Kanade, and Y. Sheikh. Convolutional pose machines. In Proceedings of the IEEE conference on Computer Vision and Pattern Recognition, pages 4724–4732, 2016.

36. K. Xu, W. Hu, J. Leskovec, and S. Jegelka. How powerful are graph neural networks? arXiv preprint 1810.00826, 2018.

37. K. Xu, C. Li, Y. Tian, T. Sonobe, K.-i. Kawarabayashi, and S. Jegelka. Representation learning on graphs with jumping knowledge networks. In International Conference on Machine Learning, pages 5453–5462. PMLR, 2018.

